# Modular Integration of Impedance Sensing for Real-Time Assessment of Barrier Integrity

**DOI:** 10.64898/2026.03.08.703312

**Authors:** Sami Farajollahi, Mehran Mansouri, Dinindu De Silva, Meng-Chun Hsu, Kaihua Chen, Aidan Hughes, Poorya Esmaili, Krittika Goyal, Steven W. Day, James L. McGrath, Vinay V. Abhyankar

## Abstract

Microphysiological systems (MPS) are essential for modeling tissue barriers, yet integrating electrical readouts often requires permanently sealed microfluidic architectures that limit access to open-well (direct-access) workflows used in bioscience laboratories. To resolve this issue, we present a modular approach in which functional components are added and removed from a standard MPS core using a magnetic interface. This design preserves compatibility with established open-well protocols for seeding and downstream analysis, while microfluidic perfusion or electrical sensing capabilities are added only when needed. We demonstrate this approach with an impedance-sensing module that enables continuous impedance measurements to assess barrier function. By fitting spectra to an equivalent circuit model, we quantify junctional and non-junctional electrical contributions to barrier integrity over time, alongside conventional single-frequency TEER, and complementary permeability and imaging readouts. We apply this platform across three representative use cases, including LPS-induced disruption, shear stress–mediated strengthening, and compatibility with barrier models formed above a 3D hydrogel matrix.

## 1. Introduction

Tissue barriers formed by endothelial cell layers regulate molecular and cellular exchange between blood and tissue compartments and are essential for maintaining physiological homeostasis^1^. Disruption of these monolayers contributes to vascular inflammation, neurodegeneration, and fibrosis^2–4^. To investigate how barrier function responds to defined stimuli, microphysiological systems (MPS) have emerged as versatile tools that replicate tissue-level architectures and recreate dynamic biophysical and biochemical cues to support quantitative studies of transport, signaling, and regulation^5,6^.

Barrier integrity is commonly evaluated using a complementary suite of structural, transport, and electrical assays^7,8^. Fluorescence imaging visualizes the continuity of junctional protein complexes connecting adjacent cells, permeability assays quantify molecule transport across the monolayer, and electrical measurements provide quantitative, non-destructive readouts of the barrier tightness^9,10^. These assays are routinely performed in Transwell-style formats, which are open-well (direct-access), two-compartment barrier models with established protocols for seeding, stimulation, imaging, and electrical measurements. Imaging and transport measurements are widely accessible in MPS platforms, but electrical readouts remain less common because their implementation often relies on embedded electrodes integrated into permanently sealed microfluidic systems designed to prevent fluid leakage^11–13^. While sealed microfluidic formats provide unique experimental capabilities, they reduce compatibility with the open-well workflows widely used in the biological sciences^14,15^. To address this limitation, we adopted a modular design philosophy where removable functional modules are magnetically added to, or removed from, an open-well MPS core as needed^16,17^, thus combining engineering capabilities with bioscience workflows^18–20^.

As shown in **Figure 1A**, the open well µSiM (**micro**device featuring a **si**licon-nitride **m**embrane) core consists of a top well and a bottom microchannel separated by an ultrathin, optically transparent porous silicon nitride membrane that serves as the culture surface where the barrier is established. The internal components of the core can be modified to support different barrier configurations and assay formats^21–24^ (see **Figure S1**, *internal modularity*). In our approach, the core unit is seated within a bottom housing that provides a magnetic interface onto which specialized modules are added and removed on demand (*external modularity*), enabling reconfiguration between open-well and module-based formats (**Figure 1B**)^16,17,25^. Our previous work demonstrated external modular approaches, where cells were first seeded in the open-well configuration and a microfluidic flow module was then added to apply physiological shear stress, introduce neutrophils, and monitor transmigration dynamics across the endothelial layer in real time^15,26^.

**Figure 1.**
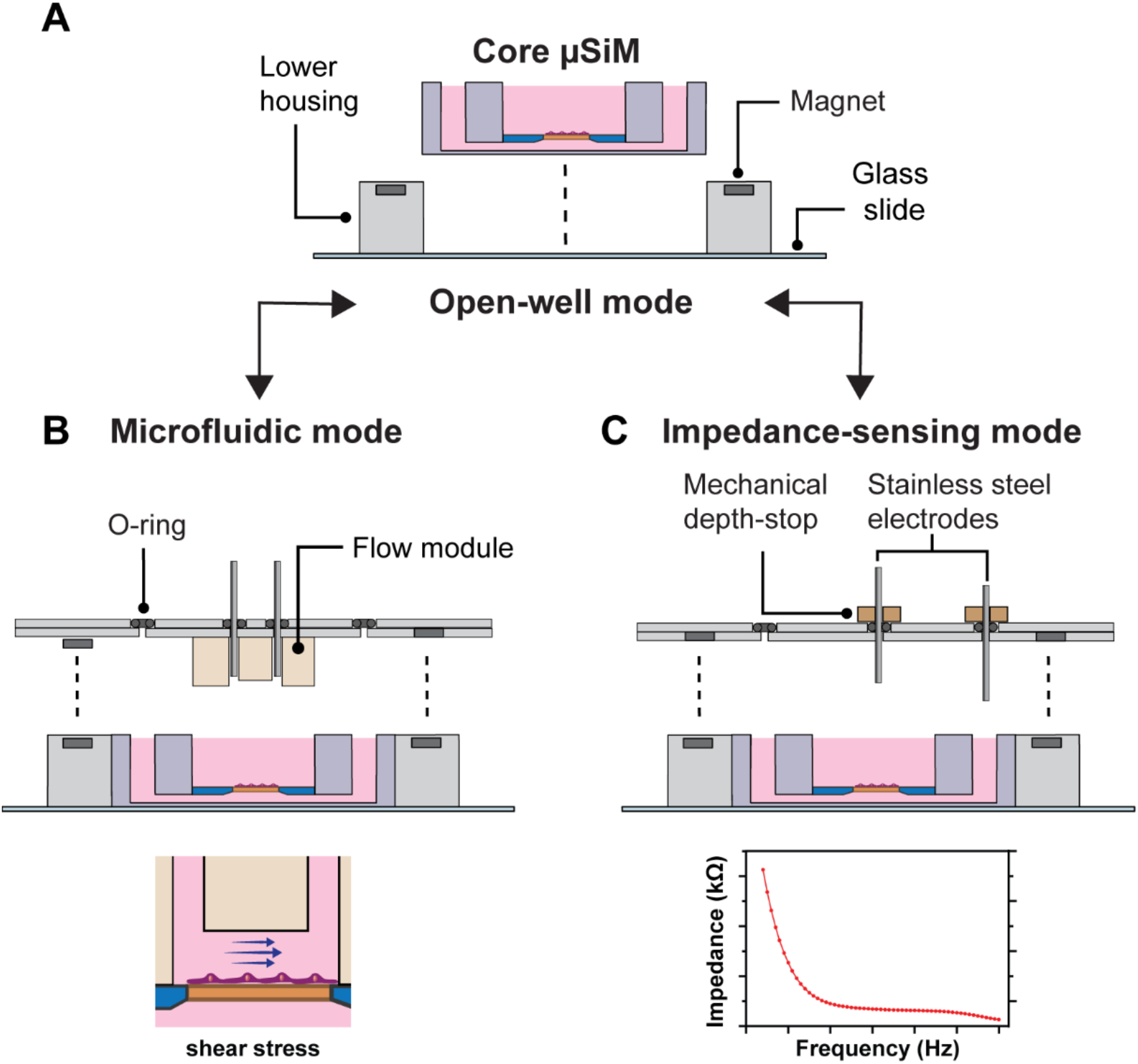
Modular reconfiguration enables on-demand microfluidic and impedance measurements with the µSiM core. **A)** Open-well mode. The core µSiM consists of a top open well and a bottom microchannel separated by an ultrathin, nanoporous silicon nitride membrane. The core is seated on a glass slide within a magnet-containing lower housing that provides the magnetic interface for reversible module attachment. **B)** Microfluidic mode. A PDMS flow module is magnetically attached to the lower housing to enable perfusion and application of shear stress. **C)** Impedance-sensing mode. An impedance module is magnetically attached to the lower housing to position stainless steel electrodes with a mechanical depth stop for repeatable placement, enabling impedance spectra collection.

We take a similar approach to introduce electrical measurement capabilities as an on-demand module. Single-frequency transendothelial electrical resistance (TEER) measurements are commonly used in Transwell-like systems and provide an accepted metric to assess barrier strength^27^. In contrast, electrical impedance spectroscopy, coupled with equivalent-circuit modeling, can resolve barrier-associated contributions from paracellular junctions, transcellular pathways, and monolayer capacitance, providing more detailed insight into properties that contribute to barrier integrity^28–31^. In this work, we demonstrate these electrical measurement capabilities across HUVEC and iPSC-derived endothelial monolayers, including hydrogel-integrated configurations. To maintain compatibility with common laboratory components, we used blunt stainless steel syringe needles as the measurement electrodes, leveraging hardware commonly used for microfluidic perfusion. We demonstrate the modular approach through three representative multi-step workflows that require device reconfiguration: 1) LPS-induced disruption, 2) shear-stress–mediated enhancement, and 3) barrier maturation above a hydrated 3D hydrogel. Across these experiments, impedance spectroscopy with equivalent circuit modeling quantifies time-varying resistive and capacitive contributions to barrier integrity and is paired with single-frequency TEER, permeability, and imaging-based readouts.

## 2. Materials and Methods

### 2.1 µSiM Platform Overview

The modular µSiM and impedance module were fabricated to enable real-time, quantitative barrier analysis while maintaining open access workflows^15^. The µSiM consists of upper and lower components manufactured at ALine Inc. (Signal Hill, CA), that sandwich an ultrathin (100 nm), nanoporous (~60 nm) silicon nitride membrane manufactured by SiMPore, Inc.^32^ (West Henrietta, NY). The nanoscale thickness and pore size approximate the structure of native basement membranes, supporting confluent endothelial or epithelial monolayers while maintaining optical transparency for imaging. The core µSiM is placed into a common bottom housing, which contains magnets. Functional modules, including flow and impedance, can be reversibly attached to this base through magnetic assembly, allowing the system to transition between open-well culture and module-based configuration as needed.

### 2.2 Impedance Module Fabrication and Assembly

#### 2.2.1 Module Housing and Magnetic Assembly

For the impedance module, 2 mm and 3.5 mm PMMA sheets were laser-cut to form the top and bottom housings, respectively, and the bottom housing was bonded to a glass coverslip using pressure-sensitive adhesive (PSA, 3M). Nickel-plated neodymium magnets (4.75 mm diameter, 0.34 kg pull force, K&J Magnetics) were press-fit into each housing to ensure self-alignment and repeatable attachment **(Figure 2)**.

**Figure 2.**
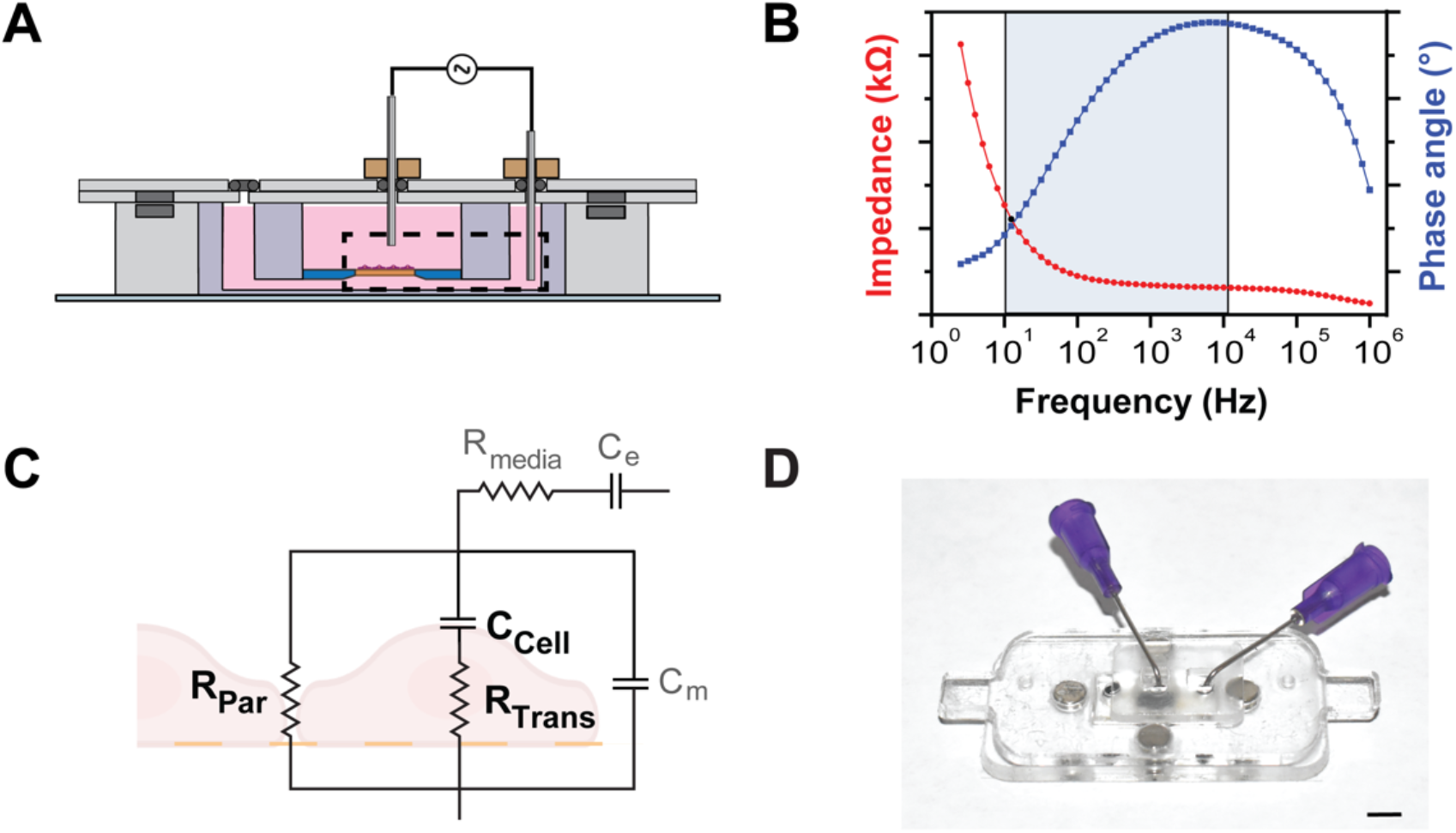
Impedance module configuration and equivalent-circuit modeling of endothelial barrier impedance. **A)** Cross-sectional schematic of the assembled impedance module showing stainless steel electrodes positioned in the top well and bottom channel compartments. The dashed box indicates the barrier region represented by the equivalent-circuit model. **B)** Representative Bode plot showing impedance magnitude (red) and phase angle (blue) across the measured frequency range; the shaded region indicates the frequency band (10 Hz–10 kHz) used for equivalent-circuit fitting. **C)**Equivalent-circuit model corresponding to the dashed region in **A**, used to extract barrier-associated resistive and capacitive parameters from impedance spectra. **D)** Photograph of the magnetically assembled device. Scale bar = 5 mm.

#### 2.2.2 Electrode Positioning and Mechanical Constraints

Polished 90°-angled 304 stainless-steel blunt syringe needles (Gauge 21, 0.82 mm O.D.) served as electrodes. Stainless steel was selected for its corrosion resistance, biocompatibility, and commercial availability. Electrode position was controlled using mechanical PDMS depth stops and O-rings within the module (**Figure 3C**). Silicone O-rings (sizes 006 and 008, McMaster-Carr) stabilized electrode positions and prevented evaporation^33^. Depth stops were adjusted to position the bottom electrode at the lowest achievable depth without contacting the channel bottom, while the top electrode was positioned in the middle of the top well. Reproducibility of electrode positioning in the module was validated via fifteen cycles of assembly, measurement, and disassembly.

**Figure 3.**
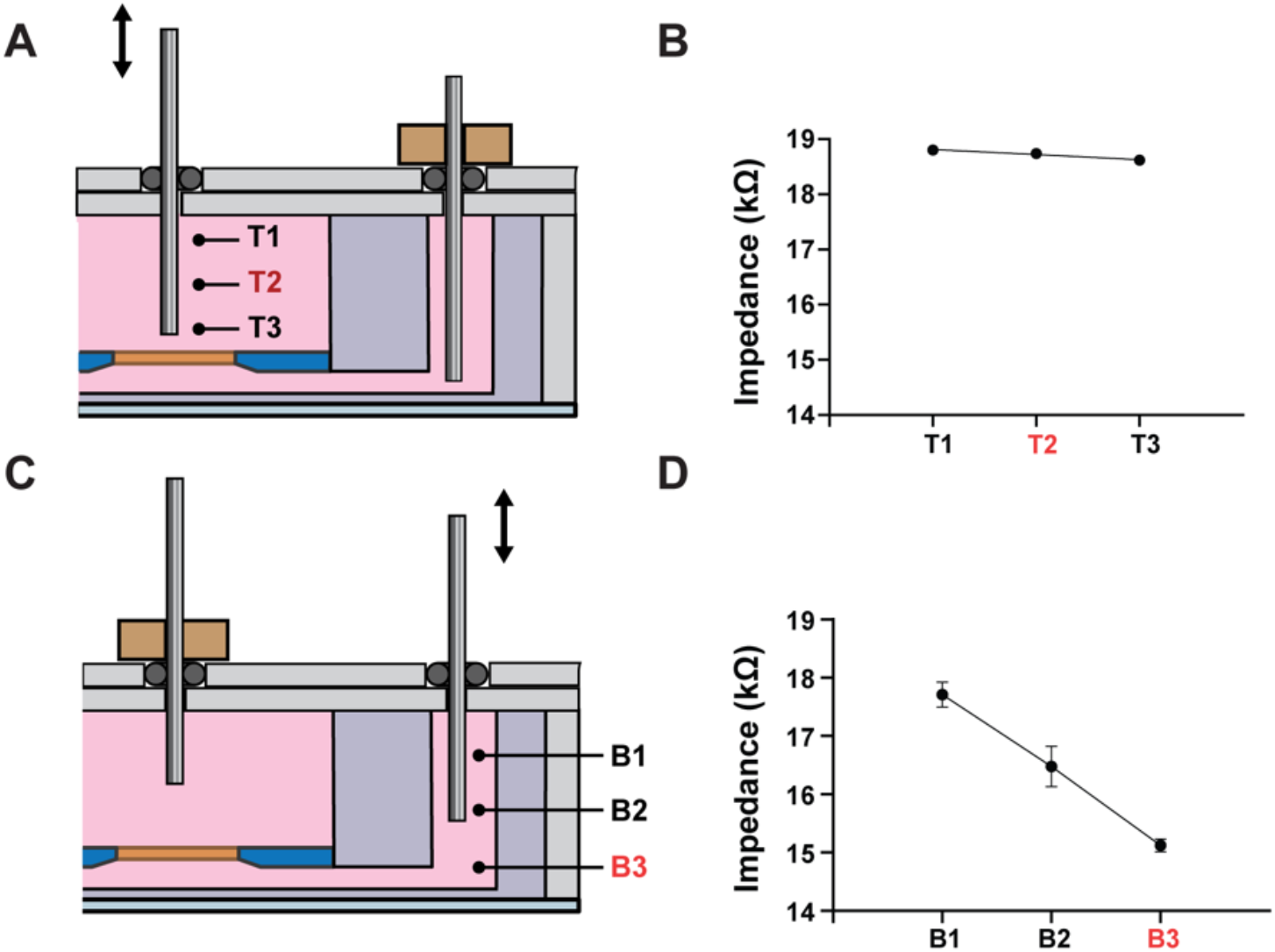
Electrode positioning to minimize background impedance. **A)** Tested positions for the electrode in the top well (T1–T3). **B)** Background impedance shows low sensitivity to top electrode position. **C)** Tested positions for the electrode in the bottom channel (B1–B3). **D)** Background impedance shows higher sensitivity to bottom electrode position. The red labels T2 and B3 indicate the final selected positions used to minimize background impedance in the µSiM core geometry.

### 2.3 Electrical Impedance Spectroscopy Measurements

#### 2.3.1 Measurement Setup and Acquisition Parameters

All components were sterilized by UV exposure for 30 min, and electrodes were rinsed in 70% ethanol. Before seeding, both compartments were filled with culture medium and equilibrated at 37 °C and 5% CO_2_ for 2 h while acquiring a baseline cell-free impedance spectrum. Impedance spectra were collected using a Gamry Reference 600 potentiostat in a two-electrode configuration, with the top electrode as the working electrode and the bottom as the reference. Impedance spectra were collected over a frequency range of 2.5 Hz to 1 MHz with seven data points per decade of frequency. For comparison to commonly used commercial measurements, TEER was reported as the impedance-derived resistance at 12.4 Hz^27^. All TEER values were normalized to the cell growth area (0.014 cm^2^) and reported as Ω cm^2^.

#### 2.3.2 Equivalent Circuit Modeling and Parameter Extraction

To resolve the contributions of different transport pathways within the endothelial barrier, impedance spectra were analyzed using an equivalent circuit model validated for endothelial monolayers^28,34^. As shown in **Figure S2**, the model decomposes total impedance into contributions from the culture: R_media_ (the resistance of the culture medium in the top well and bottom channel), C_e_ (the double-layer capacitance of the top and bottom electrodes), R_Par_ (paracellular resistance), C_Cell_ (membrane capacitance), and R_Trans,_ (transcellular resistance), and C_m_ (a model fitting term). Impedance spectra were analyzed using custom MATLAB (R2023a) scripts. Parameter estimation was performed using a bounded nonlinear least-squares optimization routine. Although spectra were collected across a full frequency range (2.5 Hz–1 MHz), circuit fitting was restricted to the 10 Hz–10 kHz region where impedance is most sensitive to barrier properties and minimally influenced by electrode polarization effects^34,35^. Fits satisfying a goodness-of-fit criterion of R^2^ > 0.95 were retained for subsequent analysis. Barrier-associated parameters (R_Par_, R_Trans_, and C_Cell_) were reported directly from the fitted model, while TEER was reported at 12.4 Hz and normalized to the culture area for direct comparison to commercial TEER measurements.

### 2.4 Finite Element Modeling

3D finite element model was developed in COMSOL Multiphysics (v5.5, AC/DC Module) to evaluate how electrode insertion depth in the µSiM influences impedance. The model included the µSiM chamber, membrane, culture medium, and stainless-steel electrodes. The PMMA components and the non-porous regions of the silicon membrane were modeled as insulators.

To account for the impedance of the porous silicon nitride membrane window without resolving the nanoscale pore geometry, the window was modeled as a homogenized effective medium. The electrical conductivity was calculated as the product of the culture medium conductivity and the membrane porosity, following the superposition principle^9^. Similarly, the conductivity of the cell monolayer was defined as a function of confluence (97%), treating the cell layer as a superposition of insulators (cells) and conductors (paracellular gaps). The electrodes were modeled with an annular cross-section.

Simulation results were used to evaluate the sensitivity of impedance measurements to electrode positioning and guided the final electrode placement used in the module design (**Figure S3**).

### 2.5 Cell Culture

#### 2.5.1 HUVEC Culture

Human umbilical vein endothelial cells (HUVECs, Lonza) were cultured in EBM-2 Basal Medium supplemented with the EGM-2 Endothelial Cell Growth Medium-2 BulletKit (Lonza) at 37 °C and 5% CO_2_. Cells were used between passages 3–6. Cells were cultured in T25 flasks, dissociated using TrypLE (Thermo Fisher Scientific) for 3 min, centrifuged at 150 × g for 5 min, and resuspended in fresh medium. µSiM membranes were coated with fibronectin (100 µg mL^−1^, Corning) for 2 h at 37 °C before seeding. HUVECs were seeded at a density of 60,000 cells cm^−2^.

#### 2.5.2 iPSC-Derived BMEC Culture

Human induced pluripotent stem cell-derived brain microvascular endothelial cells (iPSC-BMECs) were cultured in human endothelial serum free medium supplemented with B-27 (Gibco) and human fibroblast growth factor-2 (R&D Systems). Cells were used between passages 4–6. Cells were expanded in T25 flasks, dissociated using StemPro Accutase for 3 min, centrifuged at 180 × g for 5 min, and resuspended in culture medium. To enhance cell attachment, µSiM membranes were coated with a mixture of collagen IV (400 µg mL^−1^, Sigma) and fibronectin (100 µg mL^−1^, Sigma) for 2 h at 37 °C33. Cells were seeded at 60,000 cells cm^-2^ on coated membranes^36^.

### 2.6 Endothelial Barrier Conditioning and Functional Measurements

Unless otherwise noted, endothelial monolayers were established in open-well mode, converted to impedance-sensing mode by attaching the impedance module for electrical measurements, and returned to open-well mode for endpoint permeability and imaging assays.

#### 2.6.1 Inflammatory Conditioning (LPS Exposure)

Confluent HUVEC monolayers were treated with 500 ng/mL LPS (E. coli O111:B4, Sigma-Aldrich) for 24 h while impedance was recorded continuously at 30 s intervals.

#### 2.6.2 Shear Stress Conditioning

A custom flow circulation system previously described for the µSiM platform was used to apply physiological shear stress^14,15^. Prior to use, dispensing tips, flow module, housings, and flow circuit were sterilized via UV light exposure for 30 min. The system was subsequently flushed with 70% ethanol for 3 minutes followed by PBS for 10 min.

HUVEC monolayers were cultured under static conditions for four days to establish a barrier. The flow module was then magnetically attached onto the µSiM base and connected to the circulation system. Reservoirs were filled with culture medium, and flow was driven using a peristaltic pump (Langer Instruments). Based on validated COMSOL simulations of the channel geometry, a flow rate of 300 µL min^−1^ produced a uniform wall shear stress of 5 dyn cm^−2^ over the membrane region^15,36^. Continuous impedance monitoring during endothelial barrier maturation under shear stress is shown in Supplementary **Figure S4**.

#### 2.6.3 Barrier Formation above a 3D Hydrogel

To evaluate barrier formation above a biomimetic extracellular matrix, a hyaluronic acid hydrogel (HyStem® Thiol-Modified Hyaluronan Hydrogel Kit, Advanced Biomatrix) was introduced into the bottom channel. Glycosil and Extralink-Lite components were reconstituted according to the manufacturer’s protocol and mixed at a 4:1 volume ratio. The solution was injected into the bottom channel and allowed to polymerize for 90 min at 37 °C. iPSC-BMECs were then seeded onto the membrane at 60,000 cells cm^−2^ and cultured for five days before impedance and permeability measurements. Culture medium was replaced daily.

#### 2.6.4 Barrier Permeability Measurements

Permeability assays were performed using our previously established protocols for the µSiM platform^7,36^. Briefly, the medium in the top well was replaced with 100 µL of 10 kDa FITC-dextran (1 mg mL^−1^, Invitrogen), and devices were incubated for 1 h at 37 °C. Following incubation, fluid from the bottom channel was collected and transferred to a black 96-well plate. Fluorescence intensity was measured using a SpectraMax microplate reader (Molecular Devices) at excitation/emission wavelengths of 480/520 nm. Concentrations were determined from a calibration curve generated from serial dilutions of the tracer. Permeability of the endothelial layer (P_e_) was calculated based on system permeability of the devices (P_s_) and the control (P_c_) using the following equation^7^: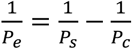.

For in situ permeability measurements in the hyaluronic acid gel, devices were placed on the stage of an Andor spinning disk confocal microscope mounted on a Nikon TiE microscope. The top well medium was replaced with Lucifer Yellow solution (150 µg mL^−1^, 457 Da, Invitrogen), and fluorescence images were acquired once per minute for 10 min. After imaging, devices were left undisturbed for 2 h to allow complete dye diffusion through the hydrogel, and a reference image was obtained.

#### 2.6.5 Junctional and Viability Imaging

Cells were fixed with 4% paraformaldehyde (Fisher Scientific) for 15 min and washed three times with PBS containing 0.05% Tween-20 (PBST). Samples were blocked with 5% goat serum in PBS for 30 min at room temperature. For VE-cadherin staining, Alexa Fluor 488-conjugated VE-cadherin antibody (Thermo Fisher, 53-1449-41) was diluted 1:100 in blocking buffer and incubated overnight at 4 °C. For Claudin-5 staining, primary antibody (Thermo Fisher, 35-2500) was diluted 1:200 in blocking buffer and incubated overnight at 4 °C. After washing, samples stained for Claudin-5 were incubated with Alexa Fluor 488-conjugated secondary antibody (Thermo Fisher, A-11001, 1:200). Nuclei were counterstained with Hoechst 33342 (1:500).

Cell viability was assessed using a LIVE/DEAD Cell Imaging Kit (Thermo Fisher, L3224) according to the manufacturer’s instructions. Quantification of viable cells was performed using CellProfiler. Imaging was performed using an Olympus IX81 inverted microscope or a Leica SP5 confocal microscope.

### 2.7 Image and Data Analysis

Image processing and quantitative analysis were performed using Fiji. Phase-contrast and fluorescence imaging was performed using an Olympus IX-81 microscope with CellSens software. Cell orientation relative to the flow direction (0°) was quantified using CellProfiler. Directionality histograms spanning 0– 90° were generated using 15° bins in GraphPad Prism (v10.4.1). Cells were considered aligned if their orientation angles fell within 0–30° of the flow direction.

### 2.8 Statistical Analysis

Data are presented as box-and-whisker plots showing the median, interquartile range (box), and full range (whiskers) from at least three biological replicates unless otherwise stated. Experimental groups and controls were conducted in parallel. Unpaired t-tests were used for two-group comparisons, and two-way ANOVA with Tukey test for time-course data. A p-value < 0.05 was considered statistically significant. Impedance measurements were performed in triplicate per condition, and circuit fits required R^2^ > 0.95. To quantify measurement consistency, the coefficient of variation (CV), defined as the ratio of the standard deviation to the mean, was used to assess reproducibility. Data analysis and plotting were performed using GraphPad Prism v10.4.1.

## 3. Results and Discussion

To provide on-demand electrical readouts within the µSiM workflow (**Figure 1A, B**), we developed a removable impedance-sensing module that magnetically attaches to the µSiM core (**Figure 1C**). Impedance spectra were fit to the equivalent-circuit model in **Figure 2** to extract R_Par_, R_Trans_, and C_Cell_, and a single-frequency TEER value was reported from the same spectra. These electrical readouts, together with complementary imaging and permeability measurements, were used to quantify barrier disruption during LPS stimulation, strengthening under shear, and barrier formation above a hydrated 3D matrix.

### 3.1 Controlled electrode positioning minimizes background impedance

Impedance measurements are sensitive to electrode position, and uncontrolled placement can introduce background variability that masks biologically meaningful changes in barrier properties^37–40^. To minimize position-related variability during module attachment, we designed the impedance module to control insertion depth and limit lateral motion using mechanical stops and integrated O-rings (**Figure 2A, B**). We then identified electrode insertion depths that minimized background impedance in the µSiM core geometry.

COMSOL simulations of the electric potential distribution (**Supporting Figure S3**) predicted a shallow impedance–depth slope for the top electrode and a steeper slope for the bottom electrode, reflecting the asymmetric field distribution between the large top reservoir and the confined bottom channel region. We tested this prediction experimentally by varying top electrode insertion depth (T1–T3) and bottom electrode insertion depth (B1–B3) (**Figure 3**). Consistent with the model prediction, background impedance varied minimally across the tested top electrode positions (**Figure 3A** and **B**), with an impedance–depth slope of −0.2 kΩ mm^−1^, while the bottom electrode showed a stronger dependence on insertion depth (**Figure 3C, D**), with a slope of −1.7 kΩ mm^−1^.

This slope difference is consistent with the underlying potential gradients in the two compartments. In the top reservoir, the electric field is diffuse and potential gradients are shallow, so small vertical shifts in the top electrode produce only minor changes in the measured background. In contrast, the bottom electrode is positioned near a confined channel region where the electric field is concentrated and the potential drops more sharply, resulting in increased sensitivity to electrode height.

Based on these results, we selected electrode positions that place the top electrode in the reservoir and away from the membrane, and the bottom electrode as close to the confined channel region as practical without risking contact with the channel floor. While the specific insertion depths that minimize background impedance depend on the device geometry, the same model-guided slope analysis provides a general approach to identify suitable electrode positions.

### 3.2 Impedance module design enables low measurement variability across multiple reconfiguration cycles

With electrode positions identified to minimize background impedance (**Figure 3**), we next asked whether the impedance module could reproducibly maintain those positions across repeated attachment and removal cycles. To quantify variability, we performed 15 reconfiguration cycles per device and collected an impedance spectrum after each attachment (Figure 4A). Impedance magnitude at 12.4 Hz derived from each spectrum remained stable across cycles for each device (**Figure 4B**), with intra-device coefficients of variation (CV) ranging from 1.2–2.4%. Because 15 reconfiguration cycles per device exceeds typical experimental needs, these measurements provide a conservative estimate of reconfiguration-associated variability, which is small relative to the biological impedance changes observed in subsequent experiments.

**Figure 4.**
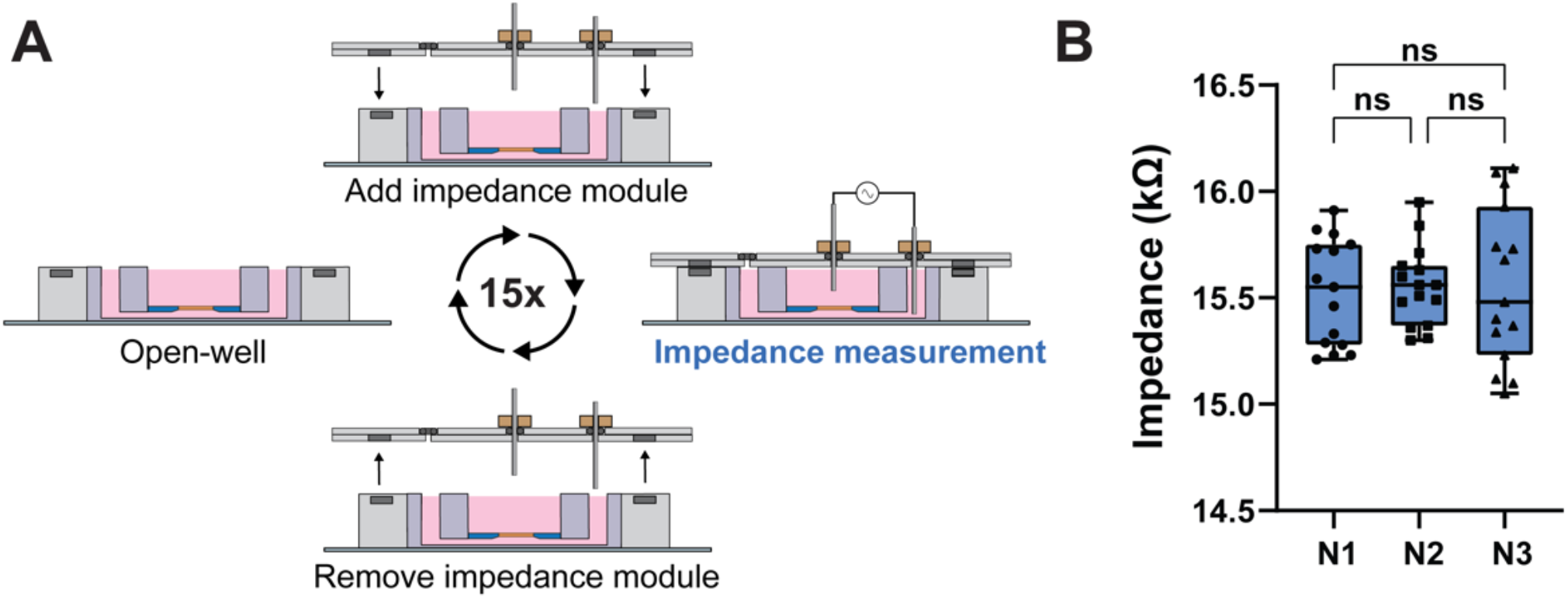
Repeatability of impedance measurements across modular reconfiguration cycles. **A)** Protocol showing repeated attachment and removal of the impedance module with impedance spectra collected after each cycle (15 cycles). **B)** Impedance magnitude at 12.4 Hz extracted from the impedance spectra across 15 cycles for three independent devices (N1–N3). Each point represents one reconfiguration cycle. Data are shown as box-and-whisker plots with median, interquartile range, and full range. N = 3 independent devices. n = 15 cycles per device. ns indicates not statistically significant.

### 3.3 Cell viability is unaffected by continuous impedance measurement

Because continuous impedance monitoring applies a sustained AC signal through stainless-steel electrodes, we next evaluated whether these measurement conditions affected cell viability. HUVEC monolayers were cultured for four days in the µSiM under static conditions before attaching the impedance module, after which impedance measurements were acquired every 30 seconds for 24 hours. LIVE/DEAD staining showed viability of 91.0% ± 1.6% for the control and 91.5% ± 3.5% for the impedance measurement condition. These results indicate that 24 hours of impedance acquisition did not negatively affect cell viability (**Figure 5A, B**). Having established compatibility with long-term culture, we next evaluated whether the impedance module could resolve biologically relevant changes in barrier properties.

**Figure 5.**
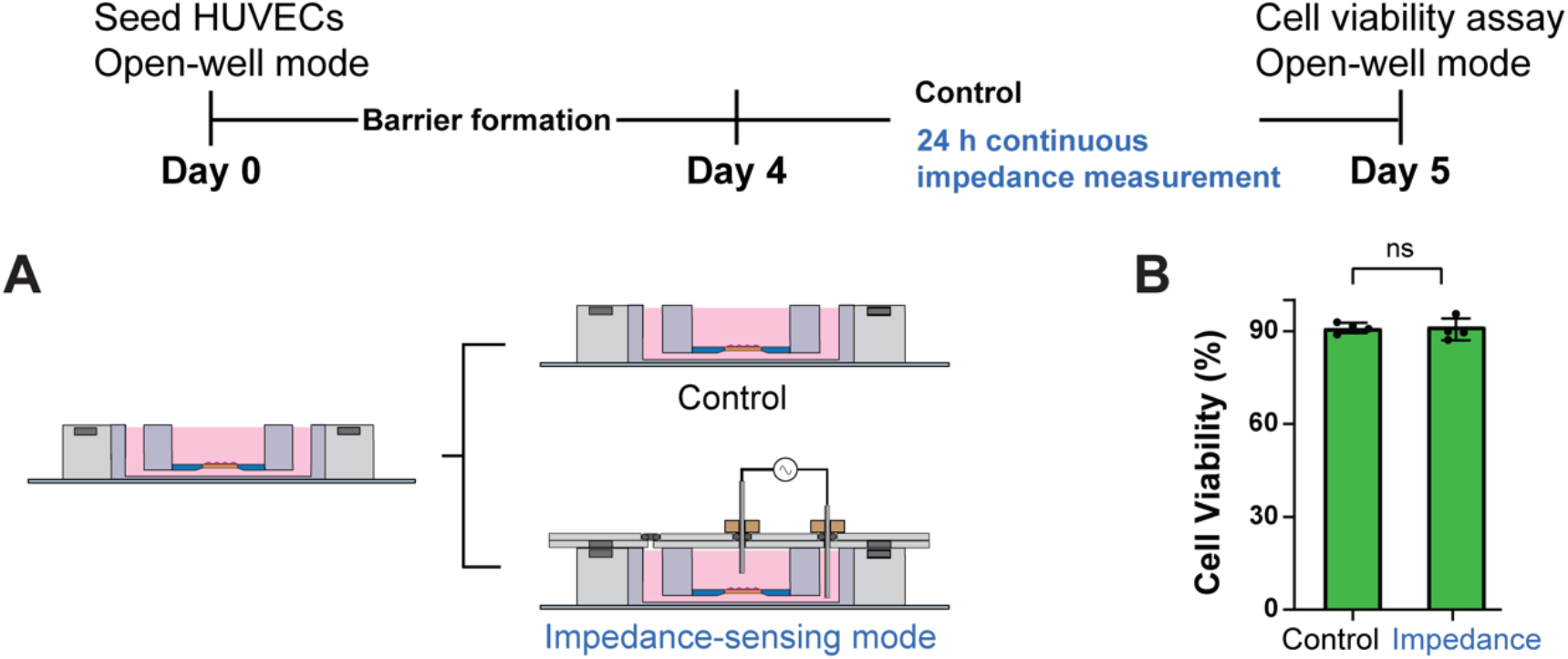
Continuous impedance measurement does not affect endothelial cell viability. **A)** Experimental workflow. HUVEC monolayers were cultured in open-well mode through Day 4, then converted to impedance-sensing mode for 24 h of continuous impedance measurement. On Day 5, LIVE/DEAD staining was performed and compared to control devices maintained in open-well mode without impedance measurement. **B)** Cell viability showed no significant difference between control (91.0% ± 1.7%) and impedance measurement (91.5% ± 3.5%) conditions. Data are shown as mean ± standard deviation. N = 4 independent devices. ns indicates not statistically significant.

### 3.4 Time-resolved measurement of LPS-induced barrier disruption

Exposure to the bacterial endotoxin lipopolysaccharide (LPS) is commonly used to disrupt endothelial barriers in vitro^41^. We used this inflammatory stimulus to test whether the modular µSiM workflow could resolve time-dependent changes in barrier integrity while preserving compatibility with endpoint assays. HUVEC monolayers were cultured for four days in the µSiM, after which LPS was added directly to the top well at t = 0. The impedance module was then attached to the bottom housing, enabling continuous impedance monitoring (**Figure 6A**).

**Figure 6.**
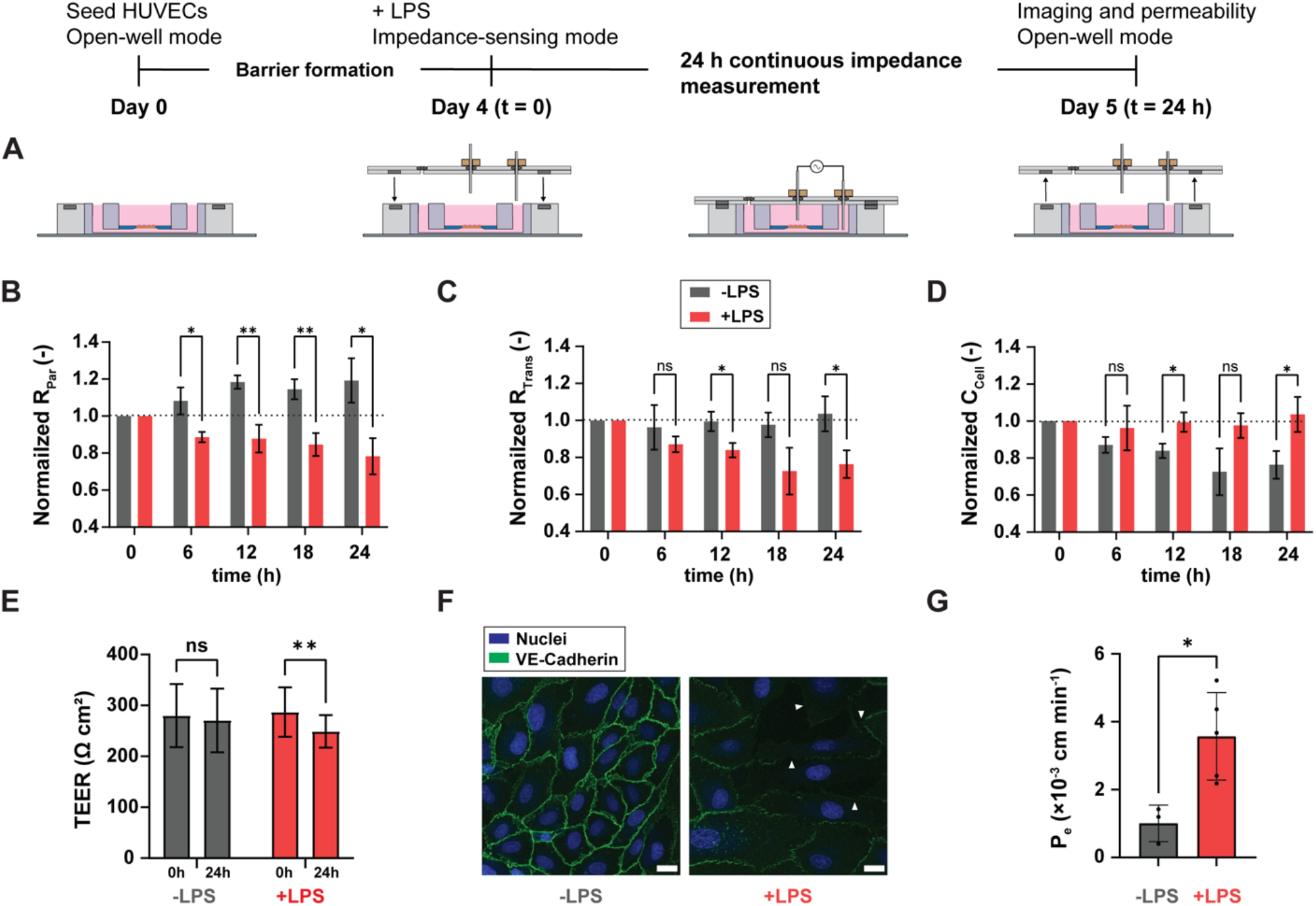
Continuous impedance measurement detects LPS-induced endothelial barrier disruption. **A)** Experimental workflow highlighting modular reconfiguration. HUVECs were cultured for 4 days in open-well mode to establish a barrier. At Day 4 (t = 0 h), LPS was added and the impedance module was attached to convert devices to impedance-sensing mode. Impedance spectra were continuously collected over 24 h. At Day 5 (t = 24 h), the impedance module was removed, and devices were returned to open-well mode for either immunofluorescence imaging or permeability assays. **B–D)** Normalized time courses of model-fit parameters obtained from equivalent-circuit analysis of impedance spectra. **E)** Single-frequency TEER at 12.4 Hz extracted from impedance spectra at t = 0 and t = 24 h for −LPS and +LPS conditions. **F)** Representative immunofluorescence images of VE-cadherin (green) and nuclei (blue) after 24 h under −LPS and +LPS conditions. Arrowheads indicate disrupted junctions in the LPS-treated monolayer. Scale bar = 20 µm. **G)** Permeability to 10 kDa dextran measured at t = 24 h. Data are shown as mean ± standard deviation. N = 3–5 independent devices per condition. *p < 0.05. **p < 0.01. ns indicates not statistically significant.

Impedance spectra fit to the equivalent circuit model revealed a time-dependent reduction in paracellular resistance with normalized R_Par_ decreasing to 0.78 ± 0.10 of its initial value over 24 hours (**Figure 6B**), with statistically significant differences observed several hours after LPS exposure. The early response was dominated by changes in paracellular resistance, and secondary responses emerged at later time points, with R_Trans_ decreasing to 0.76 ± 0.07 of baseline levels after prolonged exposure (**Figure 6C**). In contrast, C_cell_ did not exhibit an immediate effect but showed a time-dependent drift, with divergent behaviors between conditions apparent at the 24-hour point (**Figure 6D**). Together, these trends indicate that changes in paracellular resistance emerge earlier than changes in non-junctional parameters during LPS exposure.

The delayed onset of barrier weakening observed here is expected, as LPS-induced disruption depends on signaling-driven reorganization of junctional complexes rather than acute structural damage^42^. This feature makes LPS stimulation a useful assay for evaluating the temporal resolution of impedance-based measurements. For comparison to conventional metrics, TEER at 12.4 Hz decreased from 300 ± 48 to 221 ± 32 Ω cm^2^ over 24 hours in LPS-treated monolayers and remained stable in untreated controls (**Figure 6E**). While TEER reported the overall loss of barrier resistance, frequency-resolved impedance measurements combined with model-based parameter extraction resolved early and late changes in barrier properties.

Impedance-based observations were validated using complementary endpoint assays performed in compatible formats enabled by modular reconfiguration capabilities. Immunofluorescence staining showed continuous VE-cadherin junctions in control monolayers and fragmented junctions following LPS exposure (**Figure 6F**). Permeability measurements showed an approximately 3.5-fold increase in permeability in LPS-treated monolayers compared to controls (**Figure 6G**). Together, these endpoint assays confirm the barrier disruption detected during continuous impedance monitoring

### 3.5 Shear-stress–mediated barrier strengthening

Shear stress is a key physiological regulator of endothelial barrier integrity^43^, and we next evaluated whether the modular µSiM workflow could resolve shear-induced barrier strengthening. As shown in **Figure 7A**, HUVEC monolayers were first cultured to confluence in the open-well mode. On Day 4, the impedance module was attached to acquire baseline impedance spectra and replaced with the flow module. The monolayers were perfused and exposed to 5 dyn cm^−2^ shear stress for 24 hours. The flow module was then removed, and the impedance module was reattached to quantify changes in electrical barrier properties in the same samples.

**Figure 7.**
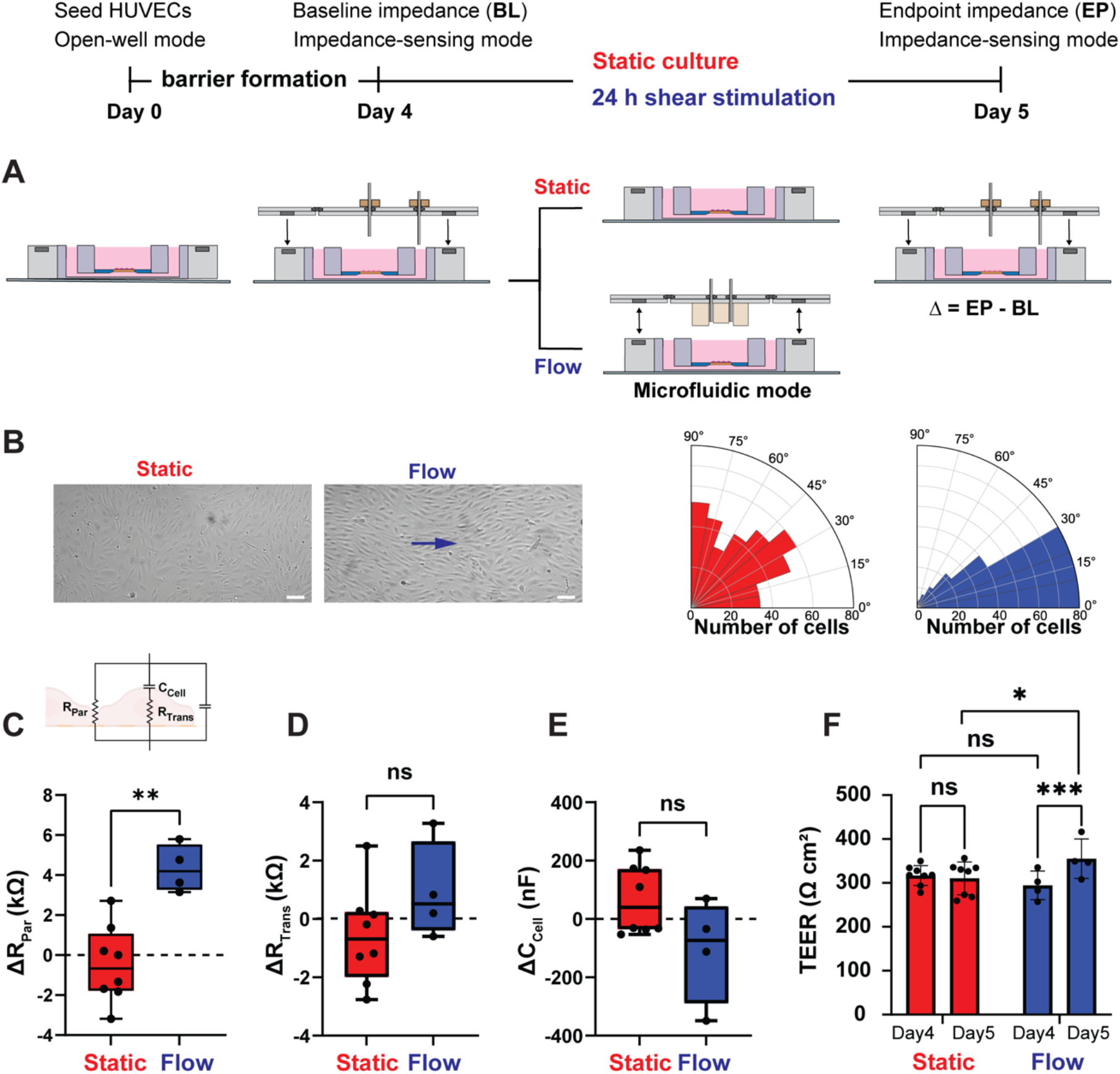
Physiological shear stress induces morphological alignment and barrier strengthening in HUVEC monolayers. **A)** Experimental workflow for parallel static and flow conditions. HUVECs were seeded and cultured in open-well mode through Day 4, then baseline impedance (BL) was measured in impedance-sensing mode. Devices were maintained for 24 h under static conditions in open-well mode or under flow in microfluidic mode to apply 5 dyn cm^−2^ shear stress. The flow module was removed, and endpoint impedance (EP) was measured at Day 5; net changes were calculated as Δ = EP − BL (Day 5 − Day 4). **B)** Representative bright-field images and polar histograms showing cell alignment under flow. Arrow indicates flow direction. Scale bar = 100 µm. **C–E)** Net changes in model-fit parameters between endpoint and baseline (Δ = EP − BL) obtained from equivalent-circuit analysis of impedance spectra. **F)** Single-frequency TEER at 12.4 Hz extracted from the impedance spectra at baseline (Day 4) and endpoint (Day 5) for static and flow conditions. Data are shown as box-and-whisker plots (median, interquartile range, full range) for (C–E) and as mean ± standard deviation for (F). N = 4–8 independent devices per condition. *p < 0.05, **p < 0.01, ***p < 0.001; ns, not significant.

HUVECs exhibited the expected morphological response to shear stress, elongating and aligning in the flow direction (**Figure 7B**). Polar histograms confirmed this reorientation, showing a shift from a random angular distribution under static conditions to pronounced alignment along the flow direction. Consistent with this remodeling, flow-conditioned monolayers exhibited a significant net increase in paracellular resistance from baseline (mean ΔRPar ≈ 4 kΩ), whereas static controls showed only minor fluctuations over the same interval (mean ΔRPar ≈ −1 kΩ) (**Figure 7C**). This difference was statistically significant (p = 0.0011), consistent with shear-mediated strengthening of junctional integrity.

In contrast to the paracellular response, net changes in transcellular resistance (ΔR_Trans_) and membrane capacitance (ΔC_Cell_) were not statistically different between the static and flow conditions (**Figure 7D–E**). These results indicate that the dominant response to shear stimulation in the monolayer arises from junctional tightening rather than changes in transcellular pathways or membrane capacitance. Although the experiments above relied on sequential reconfiguration of the flow and impedance-sensing modules, the platform also supports impedance measurements during continuous perfusion. Using a combined flow– impedance configuration (**Supplementary Figure S4**), continuous impedance showed a progressive increase in barrier resistance during multi-day shear exposure, consistent with endothelial monolayer maturation. Together, these results demonstrate that electrical sensing can be integrated alongside fluidic stimulation when required, further extending the flexibility of the modular workflow.

### 3.6 Impedance measurement of endothelial barrier formation above a hydrogel environment

Having demonstrated the ability to measure endothelial barrier disruption and strengthening in 2D configurations, we next evaluated whether impedance measurements remain compatible with barrier models that incorporate a more physiologically relevant 3D microenvironment. Microphysiological barrier models often incorporate a 3D extracellular matrix beneath endothelial monolayers to represent interfaces between vascular and tissue compartments^44,45^. We therefore tested whether impedance spectra and model-derived parameters could be resolved when the bottom channel of the µSiM was filled with a hydrogel matrix.

To model a brain-relevant barrier configuration, the bottom channel of the µSiM was filled with a hyaluronic acid (HA) hydrogel^46^ and iPSC-derived brain microvascular endothelial cells (BMECs) were cultured on the membrane (**Figure 8A**). Impedance spectra collected on Day 0 and Day 5 enabled resolution of barrier formation during culture. Paracellular resistance increased over the five-day period, with R_Par_ increasing as the endothelial barrier developed (**Figure 8B**). In contrast, transcellular resistance (R_Trans_) remained relatively constant (**Figure 8C**), and membrane capacitance (C_Cell_) increased by 174 ± 100 nF between Day 0 and Day 5 (**Figure 8D**). Consistent with the increase in R_Par_, the endpoint TEER reached a value of approximately **75 Ω cm**^**2**^ at day 5 (**Figure 8E**).

**Figure 8.**
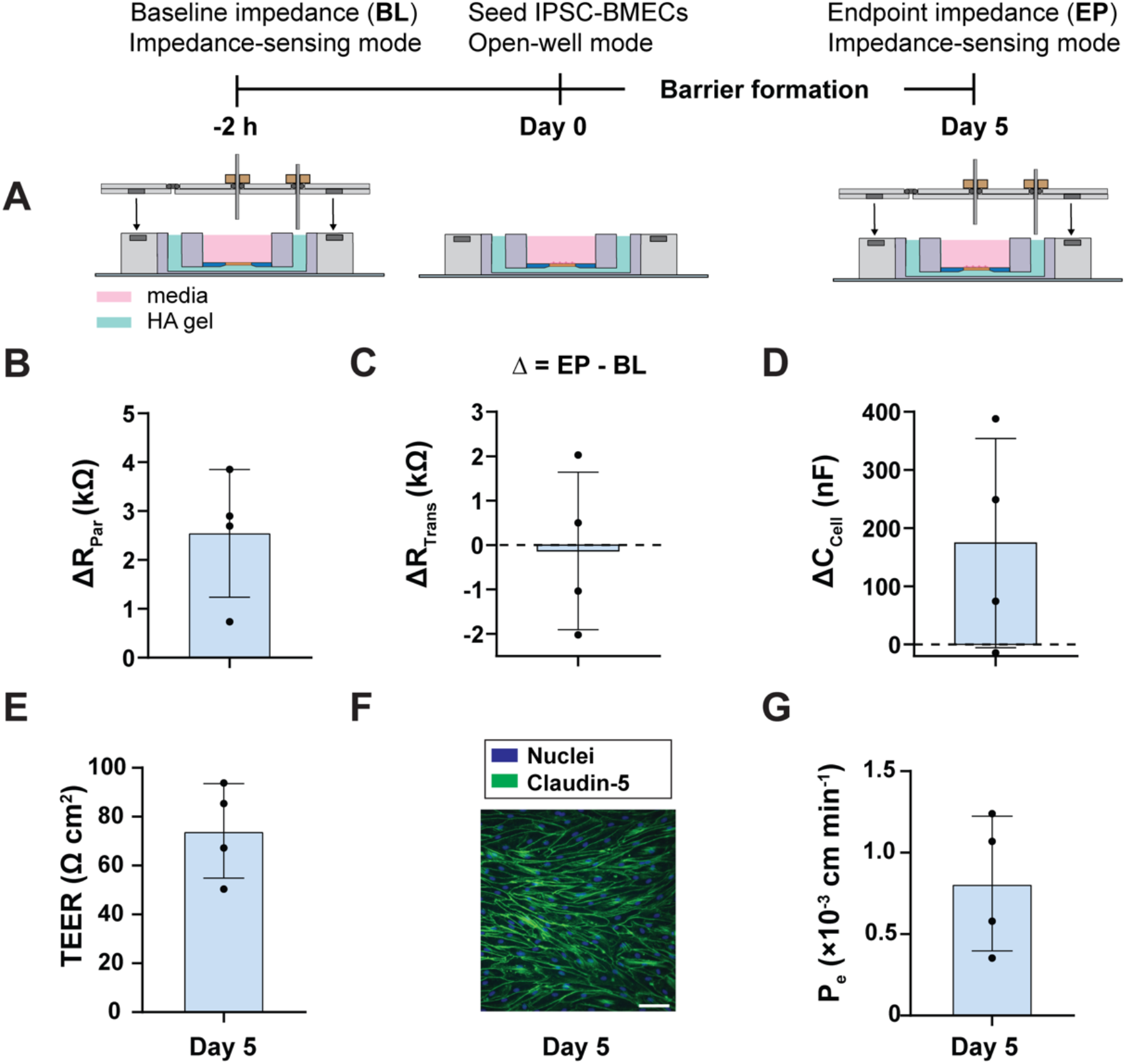
iPSC-BMEC barrier formation above a hydrated extracellular matrix. **A)** Experimental workflow and schematic of the hydrogel-integrated configuration. The bottom channel was pre-filled with a hyaluronic acid (HA) hydrogel. Baseline impedance (BL) was recorded in impedance-sensing mode on the cell-free, gel-containing device. iPSC-BMECs were seeded 2 h later in open-well mode (Day 0) and cultured to form a barrier, followed by endpoint impedance (EP) collection in impedance-sensing mode at Day 5. **B–D)** Model-fit parameters reported after subtraction of the cell-free baseline background (Δ = EP − BL) using equivalent-circuit analysis of impedance spectra. **E)** Endpoint TEER at 12.4 Hz (Day 5) extracted from the impedance spectra after subtracting the cell-free baseline background. **F)** Representative phase-contrast and immunofluorescence images at Day 5 showing Claudin-5 (green) and nuclei (blue). Scale bar = 100 µm. **G)** Endpoint permeability to Lucifer Yellow measured at Day 5. Data are shown as mean ± standard deviation; N = 4 independent devices.

Immunofluorescence staining showed BMECs formed confluent monolayers with continuous Claudin-5 expression on Day 5 (**Figure 8F**). Independent permeability measurements using a small-molecule tracer confirmed the formation of an endothelial barrier above the HA hydrogel (**Figure 8G**). *In situ* permeability values were on the order of 8 × 10^−4^ cm min^−1^, consistent with reported benchmarks for moderately tight paracellular barriers in µSiM-based blood–brain barrier models^22,47^. Together, these results demonstrate that impedance-derived barrier metrics can be resolved in the presence of a hydrated 3D matrix and are supported by complementary imaging and permeability assays, extending the modular µSiM workflow to barrier models that incorporate physiologically relevant tissue interfaces.

## 4. Conclusions

This work establishes a modular strategy for incorporating electrical measurements of barrier function in the µSiM platform. A magnetically attached impedance module provides stable, repeatable electrode positioning while preserving the open-well workflow, enabling electrical measurements to be added on demand without modifying established culture, imaging, or assay protocols.

Using this approach, impedance spectroscopy combined with equivalent circuit modeling resolved changes in barrier function arising from inflammatory challenges, shear-based conditioning, and junctional maturation in a brain microvascular model that included a hydrated extracellular matrix. The same system also provided conventional single-frequency TEER measurements in each of these experiments. These results demonstrate that electrical sensing can be integrated into microphysiological workflows as a complementary measurement modality rather than a fixed design constraint.

More broadly, the same magnetic attachment strategy supports the introduction of additional modules as experimental needs evolve, allowing measurement and control capabilities to expand without redesigning the established core device. Because these modules can be fabricated using straightforward laser cutting and lamination approaches, our work extends the modular utility of the µSiM barrier-on-a-chip platform and provides a generalizable and accessible route for integrating electrical and complementary modalities into microphysiological systems.

## 5. Acknowledgements

This research was partially supported by the NIH under award numbers U2CAG088071, R16GM146687 and R44GM137651. The authors thank Sam Lee and Allison Young from the RIT Medical Illustration MFA program for illustration support. This content is solely the responsibility of the authors and does not necessarily represent the official views of the NIH.

## References

1. Abbott, N. J., Patabendige, A. A. K., Dolman, D. E. M., Yusof, S. R. & Begley, D. J. Structure and function of the blood–brain barrier. Neurobiol. Dis. 37, 13–25 (2010).

2. Fließer, E., Lins, T., Berg, J. L., Kolb, M. & Kwapiszewska, G. The endothelium in lung fibrosis: a core signaling hub in disease pathogenesis? Am. J. Physiol.-Cell Physiol. 325, C2–C16 (2023).

3. Herminghaus, A. et al. A Barrier to Defend - Models of Pulmonary Barrier to Study Acute Inflammatory Diseases. Front. Immunol. 13, 895100 (2022).

4. Sweeney, M. D., Sagare, A. P. & Zlokovic, B. V. Blood–brain barrier breakdown in Alzheimer disease and other neurodegenerative disorders. Nat. Rev. Neurol. 14, 133–150 (2018).

5. Ingber, D. E. Human organs-on-chips for disease modelling, drug development and personalized medicine. Nat. Rev. Genet. 23, 467–491 (2022).

6. Ronaldson-Bouchard, K. & Vunjak-Novakovic, G. Organs-on-a-Chip: A Fast Track for Engineered Human Tissues in Drug Development. Cell Stem Cell 22, 310–324 (2018).

7. McCloskey, M. C. et al. The Modular μSiM: A Mass Produced, Rapidly Assembled, and Reconfigurable Platform for the Study of Barrier Tissue Models In Vitro. Adv. Healthc. Mater. 11, 2200804 (2022).

8. Vigh, J. P. et al. Transendothelial Electrical Resistance Measurement across the Blood–Brain Barrier: A Critical Review of Methods. Micromachines 12, 685 (2021).

9. Khire, T. S. et al. Finite element modeling to analyze TEER values across silicon nanomembranes. Biomed. Microdevices 20, 11 (2018).

10. Ugodnikov, A. et al. A Microfluidic Barrier-on-Chip Platform with Integrated Porous Membrane Cell– Substrate Impedance Spectroscopy. ACS Appl. Mater. Interfaces 17, 45398–45412 (2025).

11. Henry, O. Y. F. et al. Organs-on-chips with integrated electrodes for trans-epithelial electrical resistance (TEER) measurements of human epithelial barrier function. Lab. Chip 17, 2264–2271 (2017).

12. Malik, M., Steele, S. A., Mitra, D., Long, C. J. & Hickman, J. J. Trans-epithelial/endothelial electrical resistance (TEER): Current state of integrated TEER measurements in organ-on-a-chip devices. Curr. Opin. Biomed. Eng. 34, 100588 (2025).

13. Velasco, V., Soucy, P., Keynton, R. & Williams, S. J. A microfluidic impedance platform for real-time, in vitro characterization of endothelial cells undergoing fluid shear stress. Lab. Chip 22, 4705–4716 (2022).

14. Mansouri, M. et al. Transforming Static Barrier Tissue Models into Dynamic Microphysiological Systems. J. Vis. Exp. 66090 (2024) doi:10.3791/66090.

15. Mansouri, M. et al. The Modular µSiM Reconfigured: Integration of Microfluidic Capabilities to Study In Vitro Barrier Tissue Models under Flow. Adv. Healthc. Mater. 11, 2200802 (2022).

16. Abhyankar, V. V., Wu, M.Koh, C.-Y. & Hatch, A. V. A Reversibly Sealed, Easy Access, Modular (SEAM) Microfluidic Architecture to Establish In Vitro Tissue Interfaces. PLOS ONE 11, e0156341–20 (2016).

17. Hasan, M. R. et al. One-step fabrication of flexible nanotextured PDMS as a substrate for selective cell capture. Biomed. Phys. Eng. Express 4, (2018).

18. Joshi, I. M. et al. Microengineering 3D Collagen Matrices with Tumor-Mimetic Gradients in Fiber Alignment. Adv. Funct. Mater. 34, 2308071 (2024).

19. Ahmed, A. et al. Microengineered 3D Collagen Gels with Independently Tunable Fiber Anisotropy and Directionality. Adv. Mater. Technol. 6, 2001186 (2021).

20. Ahmed, A. et al. Local extensional flows promote long-range fiber alignment in 3D collagen hydrogels. Biofabrication 14, 035019 (2022).

21. Ajalik, R. E. et al. Human Tendon-on-a-Chip for Modeling the Myofibroblast Microenvironment in Peritendinous Fibrosis. Adv. Healthc. Mater. 14, 2403116 (2025).

22. Hudecz, D. et al. Modelling a Human Blood-Brain Barrier Co-Culture Using an Ultrathin Silicon Nitride Membrane-Based Microfluidic Device. Int. J. Mol. Sci. 24, 5624 (2023).

23. Masters, E. A. et al. Staphylococcus aureus Cell Wall Biosynthesis Modulates Bone Invasion and Osteomyelitis Pathogenesis. Front. Microbiol. 12, 723498 (2021).

24. Trempel, M. A. et al. Pericytes repair engineered defects in the basement membrane to restore barrier integrity in an in vitro model of the blood-brain barrier. Mater. Today Bio 35, 102361 (2025).

25. Hsu, M.-C. et al. A miniaturized 3D printed pressure regulator (µPR) for microfluidic cell culture applications. Sci. Rep. 12, 10769 (2022).

26. Ahmad, D., Linares, I., Pietropaoli, A., Waugh, R. E. & McGrath, J. L. Sided Stimulation of Endothelial Cells Modulates Neutrophil Trafficking in an In Vitro Sepsis Model. Adv. Healthc. Mater. 13, 2304338 (2024).

27. Srinivasan, B. et al. TEER Measurement Techniques for In Vitro Barrier Model Systems. SLAS Technol. 20, 107–126 (2015).

28. Günzel, D. et al. From TER to trans-and paracellular resistance: lessons from impedance spectroscopy. Ann. N. Y. Acad. Sci. 1257, 142–151 (2012).

29. Krug, S. M., Fromm, M. & Günzel, D. Two-Path Impedance Spectroscopy for Measuring Paracellular and Transcellular Epithelial Resistance. Biophys. J. 97, 2202–2211 (2009).

30. Szulcek, R., Bogaard, H. J. & Van Nieuw Amerongen, G.P. Electric Cell-substrate Impedance Sensing for the Quantification of Endothelial Proliferation, Barrier Function, and Motility. J. Vis. Exp. 51300 (2014) doi:10.3791/51300.

31. Wegener, J., Keese, C. R. & Giaever, I. Electric Cell–Substrate Impedance Sensing (ECIS) as a Noninvasive Means to Monitor the Kinetics of Cell Spreading to Artificial Surfaces. Exp. Cell Res. 259, 158–166 (2000).

32. Striemer, C. C., Gaborski, T. R., McGrath, J. L. & Fauchet, P. M. Charge-and size-based separation of macromolecules using ultrathin silicon membranes. Nature 445, 749–753 (2007).

33. Williams, M. J. et al. A Low-Cost, Rapidly Integrated Debubbler (RID) Module for Microfluidic Cell Culture Applications. Micromachines 10, 360 (2019).

34. Robilliard, L. D. et al. The Importance of Multifrequency Impedance Sensing of Endothelial Barrier Formation Using ECIS Technology for the Generation of a Strong and Durable Paracellular Barrier. Biosensors 8, 64 (2018).

35. Krug, S. M., Fromm, M. & Günzel, D. Two-Path Impedance Spectroscopy for Measuring Paracellular and Transcellular Epithelial Resistance. Biophys. J. 97, 2202–2211 (2009).

36. Chen, K. et al. Shear Conditioning Promotes Microvascular Endothelial Barrier Resilience in a Human BBB-on-a-Chip Model of Systemic Inflammation Leading to Astrogliosis. Adv. Sci. 12, (2025).

37. Lozano, A., Rosell, J. & Pallas-Areny, R. Errors in prolonged electrical impedance measurements due to electrode repositioning and postural changes. Physiol. Meas. 16, 121–130 (1995).

38. Sanchez, B., Pacheck, A. & Rutkove, S. B. Guidelines to electrode positioning for human and animal electrical impedance myography research. Sci. Rep. 6, 32615 (2016).

39. Yeste, J., Illa, X., Alvarez, M. & Villa, R. Engineering and monitoring cellular barrier models. J. Biol. Eng. 12, 18 (2018).

40. Yeste, J. et al. Geometric correction factor for transepithelial electrical resistance measurements in transwell and microfluidic cell cultures. J. Phys. Appl. Phys. 49, 375401 (2016).

41. Chan, Y. H., Harith, H. H., Israf, D. A. & Tham, C. L. Differential Regulation of LPS-Mediated VE-Cadherin Disruption in Human Endothelial Cells and the Underlying Signaling Pathways: A Mini Review. Front. Cell Dev. Biol. 7, 280 (2020).

42. Chen, Y. et al. hnRNPA2/B1 Ameliorates LPS-Induced Endothelial Injury through NF-κ B Pathway and VE-Cadherin/ β-Catenin Signaling Modulation In Vitro. Mediators Inflamm. 2020, 1–11 (2020).

43. Conway, D. E. et al. Fluid shear stress on endothelial cells modulates mechanical tension across VE-cadherin and PECAM-1. Curr. Biol. 23, 1024–1030 (2013).

44. Campisi, M. et al. 3D self-organized microvascular model of the human blood-brain barrier with endothelial cells, pericytes and astrocytes. Biomaterials 180, 117–129 (2018).

45. Floryanzia, S. D. & Nance, E. Applications and Considerations for Microfluidic Systems To Model the Blood–Brain Barrier. ACS Appl. Bio Mater. 6, 3617–3632 (2023).

46. Samanta, S., Ylä-Outinen, L., Rangasami, V. K., Narkilahti, S. & Oommen, O. P. Bidirectional cell-matrix interaction dictates neuronal network formation in a brain-mimetic 3D scaffold. Acta Biomater. 140, 314–323 (2022).

47. Nishihara, H. et al. Advancing human induced pluripotent stem cell-derived blood-brain barrier models for studying immune cell interactions. FASEB J. 34, 16693–16715 (2020).

