## Supplemental figures S1-S4 for "Modular Integration of Impedance Sensing for Real-Time Assessment of Barrier Integrity"

### Supplementary Figures S1-S4

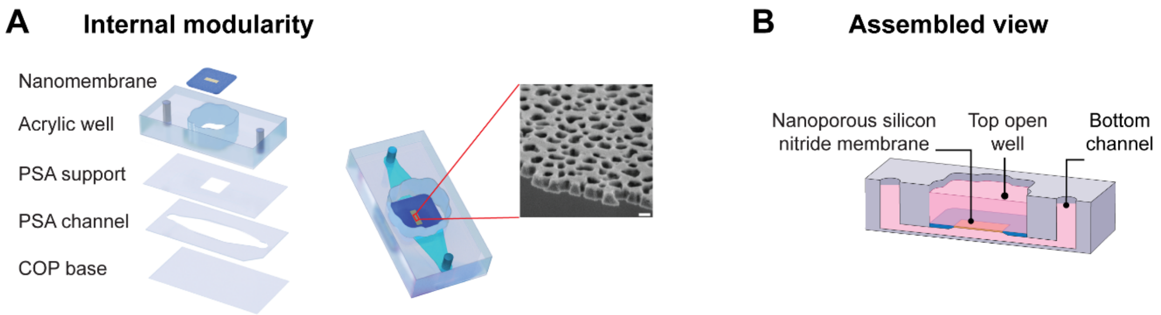


**Figure S1. Internal modularity and assembly of the µSiM platform. (A)** Exploded 3D schematic detailing the individual fabrication layers of the µSiM core, including the COP base, pressure-sensitive adhesive (PSA) microchannel and support layers, acrylic well, and the silicon nitride nanomembrane. The inset displays a scanning electron micrograph (SEM) of the nanoporous membrane. Scale bar = 100 nm **(B)** Cross-sectional schematic of the fully assembled device, illustrating the separation of the top open well and bottom microchannel by the ultrathin nanomembrane. Figure from reference 15 (Mansouri et al, 2022) reproduced with permission from Wiley.

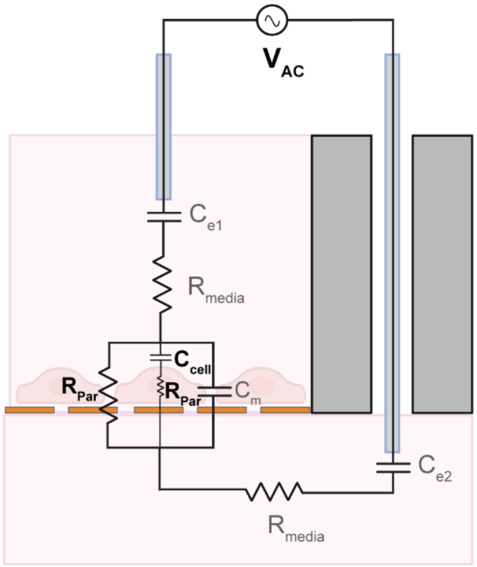


**Figure S2. Detailed equivalent circuit model of the impedance sensing setup.** Schematic representation of the complete equivalent circuit overlaid onto the cross-section of the µSiM device. The circuit maps the physical and biological components of the system to electrical elements, illustrating the electrical pathway between the top and bottom electrodes ($V_{AC}$). The model accounts for electrode double-layer capacitances ($C_{e1}$, $C_{e2}$) and bulk media resistance ($R_{media}$). The cellular monolayer is modeled by the paracellular resistance ($R_{Par}$), transcellular resistance ($R_{Trans}$), cell membrane capacitance ($C_{cell}$), and a model fitting capacitance term ($C_m$).

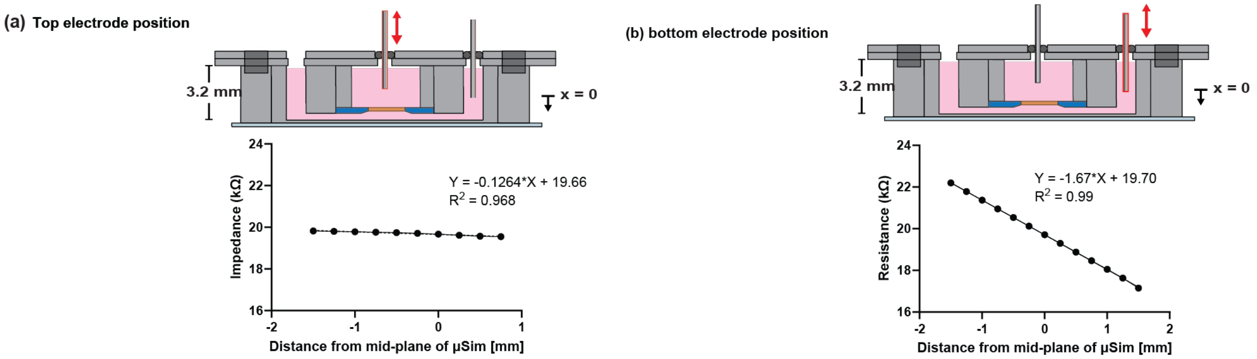


**Figure S3. Finite Element Modeling (FEM) of impedance sensitivity to electrode position.** Computational simulations were performed to guide the design of the impedance module by identifying critical geometric parameters. The model systematically varied the vertical position of the top and bottom electrodes relative to the mid-plane of the µSiM (x=0). (A) The simulation predicts that the impedance measurement is relatively insensitive to the vertical position of the top electrode, with the calculated impedance changing by only 0.13 kΩ mm-1. (B) In contrast, the simulation predicts that the measurement is highly sensitive to the vertical position of the bottom electrode, with the calculated resistance changing linearly by 1.67 kΩ/mm. These modeling results provided the core engineering rationale for the final module design, highlighting the critical need for a fixed and reproducible bottom electrode position to ensure measurement fidelity.

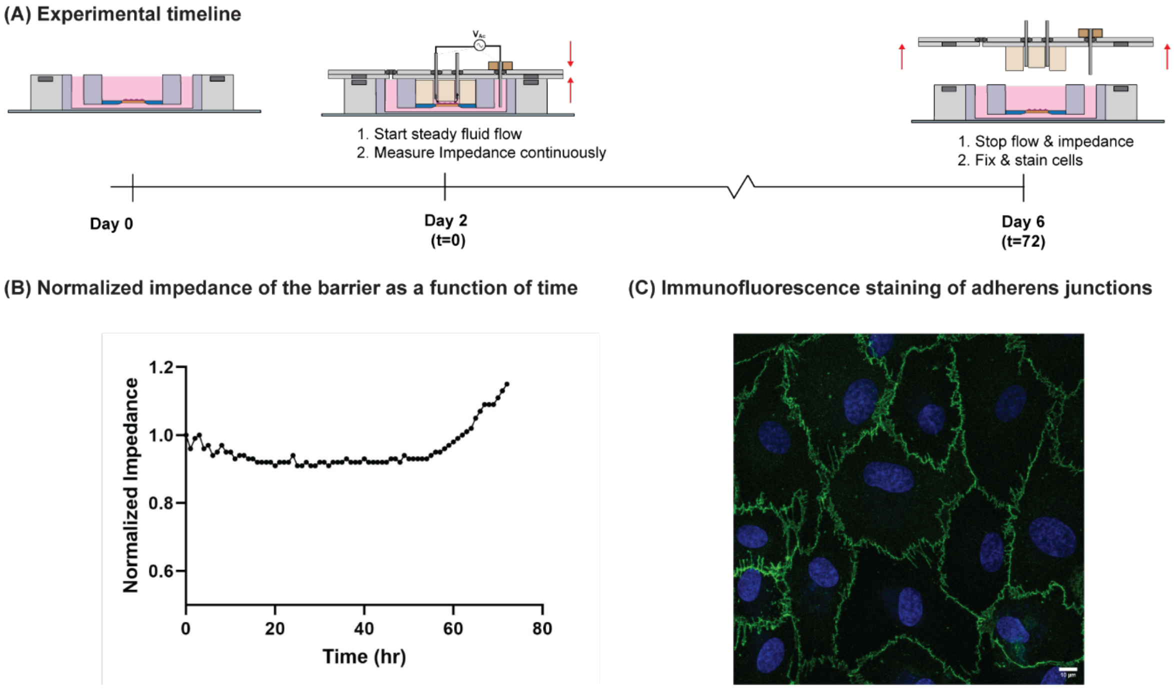


**Supplementary Figure S4. Continuous impedance monitoring of HUVEC monolayers under physiological shear stress. (A)** Experimental timeline illustrating the integration of continuous electrical monitoring with long-term fluid flow. HUVECs were seeded in the open-well µSiM on Day 0. On Day 2 (t=0), the system was reconfigured with the flow module to initiate shear stress, and continuous impedance measurements were acquired for 72 hours. On Day 6, flow was halted, and the monolayer was fixed for analysis. **(B)** Temporal profile of normalized barrier impedance over the 72-hour perfusion period. The trace reveals the dynamic response of the endothelial barrier to mechanical stimulation, demonstrating the system's stability for long-term, real-time monitoring. **(C)** Representative immunofluorescence micrograph of the endothelial monolayer after 72 hours of continuous flow and impedance recording. Staining for VE-cadherin (green) and nuclei (blue) confirms the preservation of a confluent monolayer with intact adherens junctions. Scale bar: 10 µm.
